# The human claustrum is functionally connected to cognitive networks and involved in cognitive control

**DOI:** 10.1101/461319

**Authors:** Samuel R. Krimmel, Michael G. White, Matthew H. Panicker, Frederick S. Barrett, Brian N. Mathur, David A. Seminowicz

## Abstract

The claustrum is among the most highly connected structures in the mammalian brain. However, the function of the claustrum is unknown, which is due to its peculiar anatomical arrangement. Here, we use resting state and task functional magnetic resonance imaging (fMRI) to elucidate claustrum function in human subjects. We first describe a method to reveal claustrum signal with no linear relationship with adjacent regions. We applied this approach to resting state functional connectivity (RSFC) analysis of the claustrum at high resolution (1.5 mm isotropic voxels) using a 7T dataset (n=20) and a separate 3T dataset for replication (n=35). We then assessed claustrum activation during performance of a cognitive task, the multi-source interference task, at 3T (n=33). Extensive functional connectivity was observed between claustrum and cortical regions associated with cognitive control, including anterior cingulate, prefrontal and parietal cortices. Cognitive task performance was associated with widespread activation and deactivation that overlapped with the cortical areas showing functional connectivity to the claustrum. Furthermore, the claustrum was significantly activated at the onset of the difficult condition of the task, but not during the remainder of the difficult condition. These data suggest that the claustrum can be functionally isolated with fMRI, and that it is involved in cognitive control in humans independent of sensorimotor processing.

**Highlights:**

- Removing signal from neighboring structures isolates claustrum BOLD signal at 7T and 3T field strength
- Claustrum is extensively functionally connected with cortex, including cognitive networks
- Claustrum is activated at the onset of a cognitive conflict task
- Claustrum may be involved in cognition independent of sensorimotor processing

## Introduction

In its mediolateral dimension, the claustrum is thin (submillimeter at certain points), but its rostrocaudal and dorsoventral dimensions are roughly equivalent to that of the striatum. Decades of tract tracing studies in several mammalian species indicate that the claustrum is bidirectionally connected with many cortical areas (Edelstein and Denaro, 2004; Crick and Kock, 2005; Mathur et al., 2009; Mathur, 2014; White et al., 2017; Wang et al., 2017) and is estimated by volume to be among the most highly connected structures in the brain (Torgerson et al., 2015). These observations have fueled several hypotheses that the claustrum: 1) binds multimodal sensory information for the generation of conscious perception (Crick and Koch, 2005); 2) coordinates somatosensory and motor cortical information (Smith et al., 2012) and; 3) acts as a cortico-cortical relay center supporting attention (Mathur, 2014).

Recent comprehensive analyses in a single species of how the cortical mantle connects with the claustrum demonstrate that the claustrum weakly innervates primary sensorimotor cortices, while heavily innervating frontal cortices including the anterior cingulate cortex (ACC) and the prelimbic area of the medial prefrontal cortex (PFC) (White et al., 2017). A strong input from the ACC to the claustrum also exists in rats (Smith and Alloway, 2010; White et al., 2017) and mice, and this input encodes a top-down preparatory signal that is proportional to task difficulty (White et al., 2018). These findings suggest that the claustrum may subserve frontal cortical function, including top-down executive processes (Mathur, 2014; White and Mathur, 2018). However, evidence for a role of the human claustrum supporting any of the aforementioned functional hypotheses, including cognitive processing, is particularly lacking. While the anatomical boundaries of the human claustrum can be resolved with relative ease using high-resolution structural magnetic resonance imaging (MRI), functionally resolving this structure for analysis of blood-oxygenation level-dependent (BOLD) signal with functional MRI (fMRI) is challenging, as the |signal extracted from the claustrum is heavily mixed with the signal from the neighboring insula and putamen using standard methods. Analysis of BOLD data using standard methods results in similar patterns of functional connectivity (correlation of signal between regions) when comparing claustrum, insula, and putamen. This contrasts with data from multiple tract tracing studies, which instead show unique patterns of anatomical connectivity across these regions (Nakashima et al., 2000; Mathur et al., 2009; Pan et al., 2010; Sato et al., 2013; Wang et al., 2017). Thus, standard fMRI analyses are not capable of functionally resolving the claustrum and may yield inaccurate functional connectivity and activation results.

In an effort to elucidate claustrum function, the current study has three goals: 1) devise new fMRI methodology called Small Region Confound Correction (SRCC) to functionally distinguish the claustrum from the insula and putamen, creating a corrected claustrum timeseries; 2) perform resting state functional connectivity analyses with the corrected claustrum timeseries to reveal functional coupling of the claustrum in humans; and 3) test the hypothesis that the claustrum, owing to strong connectivity with frontal cortices and recent data in mice suggesting its involvement in top-down cognitive processing (White et al., 2018), is activated during a cognitive conflict task. Our data indicate that extraction of unique claustrum signal using fMRI is possible and that the claustrum is functionally connected with cognitive networks in the resting state. During task performance, we found the claustrum to be active at the onset of – or switch to – cognitive conflict task engagement.

## Methods

### Overview

Three datasets were analyzed. The first dataset, *7T-Rest,* is publicly available and consisted of scans from 20 healthy humans scanned with 7 Tesla (T) MRI (Gorgolewski et al., 2015). The second dataset, *3T-Rest,* consisted of scans from 35 healthy humans acquired with 3T MRI. The third dataset, *3T-Task,* used the same subjects as *3T-Rest* and consisted of scans from 33 healthy humans performing a cognitive interference task acquired with 3T MRI. In *7T-Rest,* we performed seed-based whole-brain resting state functional connectivity (RSFC) analyses using the claustrum and surrounding structures (insula/putamen) as seeds. To examine claustrum functional connectivity while controlling for the influence of insula and putamen signal, we regressed out the timeseries of insula/putamen sub-regions from the claustrum, creating a corrected claustrum signal that we then used to make a claustrum specific RSFC map. We used *3T-Rest* to replicate findings from *7T-Rest.* We applied similar methods used in resting state data to *3T-Task* in order to isolate BOLD signal from the claustrum, and analyzed this signal to determine claustrum activation during cognitive load.

### Participants and MRI data

#### 7T-Rest

Data from 22 subjects were acquired from a publically available dataset scanned on a 7T MR scanner (MAGNETOM 7T, Siemens Healthcare, Erlangen, Germany). We used data from only the first session of this dataset. These data included a T1 weighted structural 3D MP2RAGE image that was used in preprocessing (TR=5000ms, TE=2.45ms, voxels=.7 mm isotropic). *7T-Rest* also included two eyes-open resting state scans using echo planar imaging (EPI) to measure BOLD fMRI while fixating on a plus sign (whole brain coverage, TR=3000ms, TE=17ms, voxels=1.5mm isotropic, slices=70, duration=300 TR). Details of the scans can be found in (Gorgolewski et al., 2015). Two subjects were excluded, one from errors induced by preprocessing and a second from different scanning parameters than the other participants, leaving 20 subjects for analysis (10 women, average age = 25, s.d. = 2).

#### 3T-Rest

36 healthy subjects were recruited as a control sample for an ongoing clinical trial and only a baseline scan was used for the following analyses. MRI data were acquired at the University of Maryland, Baltimore Medical Imaging Facility with a Siemens 3T Tim Trio scanner with 32 channel head coil (n=22) or a Siemens 3T Prisma scanner with a 64 channel head coil (n=14) due to a scanner upgrade during data acquisition. We acquired a T1 weighted structural 3D MPRAGE scan that was used in preprocessing (whole brain coverage, TR=2300ms, TE=2.98ms, voxels=1.00 mm isotropic). We also acquired eyes-open resting state scans using EPI while subjects fixated on a plus sign (whole brain coverage, TR=2s, TE=28ms, voxels=3.4 x 3.4 x 4.0 mm, slices=40, duration=300 TR). One subject was excluded due to poor coverage leaving 35 subjects (31 women, average age=37, s.d. = 13).

#### 3T-Task

Participants in the *3T-Rest* also performed the multi-source interference task (MSIT; Bush et al., 2003) as a measure of cognitive conflict. Subjects were first trained on the MSIT outside of the scanner. During the MRI session, they performed the MSIT in two runs of about 5 minutes each while echo planar imaging with whole brain coverage was acquired (TR=2500ms, TE=30ms, voxels=3.00 mm isotropic, slices=44, duration=121 TR). The task was the same as previously reported (Seminowicz and Davis, 2007, Seminowicz et al., 2011), but is briefly described here. On each trial, the volunteer was presented with an array of three numbers. In each array, two numbers were the same and one number was different. The volunteer was instructed to press a button that corresponded to the number that was different from the two other numbers or characters presented on the screen for that given trial. The control condition was a sequential tapping task in which an asterisk appeared in the same order moving from the left to the right of the screen and the subject pressed a button corresponding to the position of the asterisk. There were two levels of task difficulty (easy, difficult), which were performed in separate 20s blocks (10 stimuli per block). In the easy condition, the different number indicated the position of that number in the array (e.g. “1-2-2”, “3-2-3”, “1-1-3”), and in the difficult condition, the different number did not indicate the position of that number in the array (e.g. “3-2-2”, “3-1-3”, “1-1-2”). Three subjects were excluded because of poor coverage and missing data, leaving a sample size of 33 (29 women, average age=37 s.d.= 12).

### Initial Data Preprocessing

Data were preprocessed in SPM12 (http://www.fil.ion.ucl.ac.uk/spm/software/spm12/), including slice timing correction, realignment (motion correction), coregistration of the T1-weighted structural scan to the mean realigned functional image, segmentation of the structural scan, normalization of the structural and realigned functional images to a standard MNI template, and smoothing with a 6mm full width at half maximum (FWHM) Gaussian kernel for *7T-Rest, 3T-Rest,* and *3T-Task.* We elected to use the same smoothing kernel for 3T-Rest and 7T-Rest to have as comparable processing pipelines as possible. We did not observe obvious qualitative differences for smoothing *7T-Rest* at 6mm FWHM vs a 3mm FWHM kernel (S1).

### Analysis of resting state claustrum connectivity

#### Resting-state preprocessing

Whole-brain *7T-Rest* and *3T-Rest* data underwent the following further preprocessing before functional connectivity analyses were performed. Resting-state preprocessing and seed-based analyses were conducted in the Conn toolbox version 17f (http://www.nitrc.org/proiects/conn). Given continued controversy, we elected not to control for global signal in our analyses (Murphy and Fox, 2017). To control for noise present in white matter and CSF, we used aCompCor (Behzadi et al., 2007; Muschelli et al., 2014) to determine the first five eigenvectors of white matter and CSF. As we did not want to remove global signal, we used a twice eroded CSF mask and a white matter mask with four erosions (that did not include external or extreme capsules), as this level of erosion has been shown to no longer contain global signal (Power et al., 2017). We removed motion related signals estimated from the realignment parameters along with the first order derivatives of these parameters, in addition to the 10 eigenvectors from white matter and CSF. To avoid the reintroduction of noise while removing low frequency artifact, we used linear detrending and also simultaneously high pass filtered our voxelwise and regressor data with a cutoff of 0.008 Hz. No low pass filter was used because signal is present above standard cutoffs (∼0.08 Hz, Smith et al., 2013). Despiking was employed after these steps to remove any additional artifact that had not yet been removed (Petel et al., 2014).

### Standard analysis

Following artifact removal, ROI analyses were performed by extracting the average timeseries of all voxels within a given set of ROIs using normalized, but not smoothed data. For each subject in *7T-Rest,* left and right claustrum ROIs were hand drawn on the subject’s structural image, along with ROIs for the insula and putamen. Fig 1A shows an example tracing for one subject and Fig 1B shows the average ROIs on a group template. In *3T-Rest,* we used mean ROIs obtained from *7T-Rest,* after confirming that these mean images fit well onto the normalized structural data for *3T-Rest.* Functional connectivity was calculated as the Pearson correlation between the timeseries for each ROI and all voxel timeseries across the brain. In the standard analysis for *7T-Rest*, we used the un-corrected claustrum, whole insula, and whole putamen timeseries from individually drawn ROIs to compute RSFC across the whole brain.

**Fig 1.**
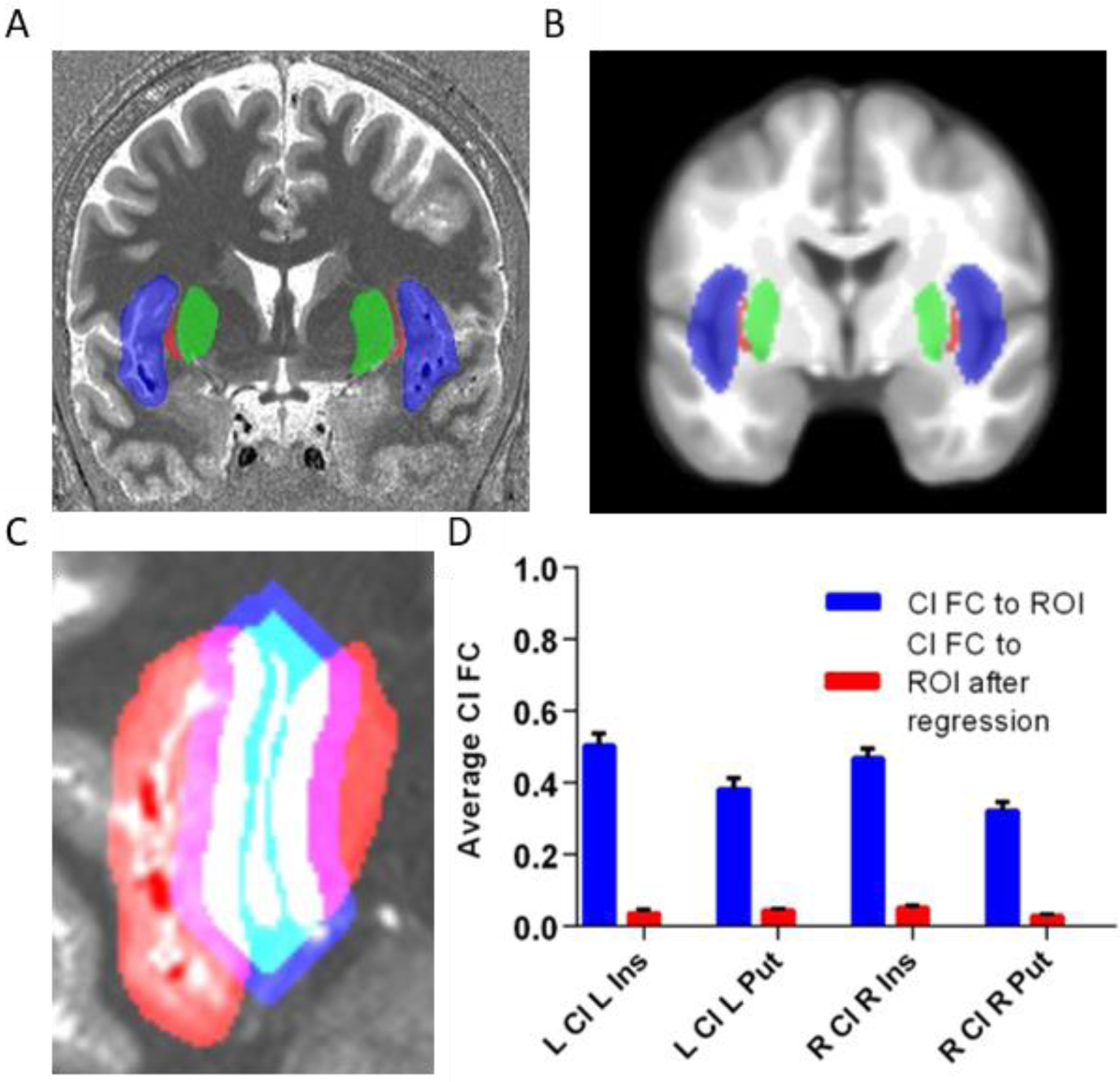
The claustrum is a thin sheet of gray matter and its fMRI signal is normally confounded by neighboring structures. A) Structural MRI image with insula (blue), claustrum (red), and putamen (green) for one subject. These regions were hand-drawn for each subject on their native anatomical image. B) Average insula (blue), claustrum (red), and putamen (green) averaged over 20 hand-drawn images displayed on an average template. C) To remove signal from neighboring structures, the claustrum was dilated, adding 6 mm around the claustrum, while maintaining its shape (blue). This dilated claustrum was overlapped with the neighboring insula and putamen (red). Voxels that contained insula/putamen and dilated claustrum separated by at least 3mm from the original claustrum were categorized as flanking regions. Flanking regions, shown in purple, were regressed out of the claustrum signal to create a corrected claustrum timeseries. D) Correlation (FC) of claustrum timeseries with L/R insula/putamen before and after regressing out flanking regions (i.e. following Small Region Confound Correction (SRCC)). The average connectivity between claustrum and its neighbors approaches zero following this approach. Error bars show standard error of the mean. Cl=claustrum, Ins=insula, Put=putamen, ROI=region of interest, FC=functional connectivity.

### Corrected claustrum analysis with Small Region Confound Correction (SRCC)

To remove the possible influence of insula and putamen signals on claustrum timeseries and hence FC, we determined the timeseries of parts of the insula and putamen that ‘flanked’ the claustrum and treated these as confounding sources for the claustrum. These flanked ROIs were calculated on the individual level for *7T-Rest* by dilating the claustrum 4 functional voxels (6 mm), and determining the overlap between the dilated claustrum and the insula/putamen seeds that were at least 2 functional voxels (3 mm) separated from the original claustrum. This resulted in ‘flanking’ regions that were continuous with insula and putamen and similar in shape to the claustrum, but still distant enough from the claustrum to ensure that they contained no claustrum data within the seed, as shown in Fig 1C. We call this approach Small Region Confound Correction (SRCC), as it is designed to eliminate partial volume effects that are particularly problematic in small regions of the brain. In *3T-Rest,* this process was repeated, but for the mean ROIs only. We regressed each claustrum timeseries (left and right) on the timeseries from the ipsilateral ‘flanking’ regions, along with previously described artifacts (motion, CSF, etc.), and the residuals from this analysis constituted a corrected claustrum timeseries (e.g. corrected for signal in the flanking insula and putamen sources) that was then used as the seed in whole-brain seed-to-voxel analysis.

To determine statistical significance of seed-to-voxel functional connectivity maps, we performed one sample t-tests in SPM12 for each RSFC map (left and right corrected claustrum). These outputs were used for qualitative comparisons between regions. A cluster-forming threshold of p<0.001 was used for all analyses and significant clusters based on FWE correction are reported, as these thresholds have been shown to adequately control for false positive rates (Woo et al., 2014).

### Modelling task data

We determined the reaction time (RT) for every trial presented in the MSIT. If the participant did not respond to a particular trial, we designated the RT for that trial as the maximum possible trial duration (i.e. 1500ms). We did not analyze accuracy, as the average was over 90 percent. Typically, the MSIT is modelled in a blocked design, using three types of blocks (tapping, easy, and difficult). However, based on observations of behavioral performance suggesting that the first one to three trials had much poorer performance (see Results) – likely reflecting the cognitive adaptations (or switching) to a new set of rules – we modelled onset and block separately, using an event design to model task onset and a block design to model the remainder of the task.

### Preprocessing

For whole brain task analyses, we did not treat insula and putamen signals as a source of noise. For analyses of claustrum activation we used similar approaches as described in *Analysis of resting state claustrum connectivity* and *corrected claustrum analysis with SRCC* (see above) except we did not include despiking nor the first derivative of realignment parameters. This approach created a corrected claustrum timeseries during MSIT that was then used to determine claustrum activation during the task. We then estimated whole brain and corrected claustrum ROI activation patterns.

### Analysis

To determine significant task activation for whole brain analyses, we performed one sample t-tests for contrast maps with a cluster-forming threshold of p<0.001 and a FWE cluster correction. Contrast maps for difficult and easy blocks and onsets were calculated based on a tapping baseline (e.g. [difficult > tapping] and [easy > tapping]). To determine claustrum activation, we performed a one-sample t-test on extracted activation estimates for each contrast extracted using MarsBaR v 0.44 (http://marsbar.sourceforge.net/). To examine the overlap between claustrum connectivity and task-related activation during MSIT, we created an overlap map between areas of significant deactivation/activation and areas with significant claustrum functional connectivity. We then used this overlap map to calculate the percentage of task-responding voxels that functionally connected with the claustrum. Average RT for each task condition (tapping, easy, difficult) was calculated for each subject, and these subject-averaged RTs were compared between each task type using a one-way repeated measures ANOVA with Greenhouse-Geisser correction. Post-hoc t-tests were used to identify specific task type differences with Tukey’s correction for multiple comparisons. We also performed a one-way ANOVA with Greenhouse-Geisser correction and used a Dunnett’s test to compare the first trial to all other trials. Whole brain contrast of difficult vs easy tasks gave task-positive (difficult>easy, EMN) and task-negative (easy>difficult, DMN) networks. For event-related plots in Fig 4C, we extracted timeseries data for the claustrum and for EMN and DMN. These timeseries were each averaged for the task of interest.

## Results

### Resting state connectivity of the claustrum at 7T: methodological approach

We used high spatial resolution fMRI data in *7T-Rest* and hand drawn ROIs of claustrum, insula, and putamen to analyze whole brain functional connectivity of these ROIs. Despite excellent resolution, we found that functional connectivity between the claustrum and the insula/putamen in this standard analysis was high (average FC of *L/R* claustrum with insula/putamen ranged from 0.32 to 0.5; Fig 1D). This contrasts with known unique connectivity of the claustrum relative to insula and putamen (Nakashima et al., 2000; Mathur et al., 2009; Pan et al., 2010; Sato et al., 2013; Wang et al., 2017), and suggests that even with high spatial resolution and well identified ROIs that insula and putamen signals are sampled in claustrum voxels. This confounds the interpretation of small structure studies in fMRI even when the regions can be identified with relative ease in structural scans.

### Creating a corrected claustrum signal with SRCC

To mitigate the effect of insula/putamen sampling in claustrum voxels, we generated a corrected claustrum timeseries using SRCC by regressing out signal from insula and putamen regions separated by two voxels from claustrum (3 mm) as sources of noise, identical to how nuisance white matter (WM) and cerebro-spinal fluid (CSF) signals are treated (see Fig 1C). This approach eliminated the linear relationship between claustrum and the surrounding structures, and the resulting average correlation between the corrected claustrum timeseries and neighboring regions reduced effectively to zero (Fig 1D).

### Resting state connectivity of the corrected claustrum at 7T and 3T

We estimated functional connectivity of the corrected claustrum timeseries across the whole brain in two datasets. Given previous findings in rats (White et al., 2017), we anticipated claustrum RSFC to be widespread and to feature connectivity with cingulate cortex, PFC, visual cortex, and intraparietal sulcus (IPS), among other regions. In *7T-Rest,* this analysis identified connectivity with the: thalamus, particularly the pulvinar; nucleus accumbens; visual cortex; both anterior and posterior cingulate cortex; PFC (including the dorsal lateral PFC, medial PFC, and ventral lateral PFC); precuneus; angular gyrus; sensorimotor cortex; parahippocampal gyrus; superior and inferior temporal gyri; and IPS (Fig 2A and B and Table S1 and S2).

**Fig 2.**
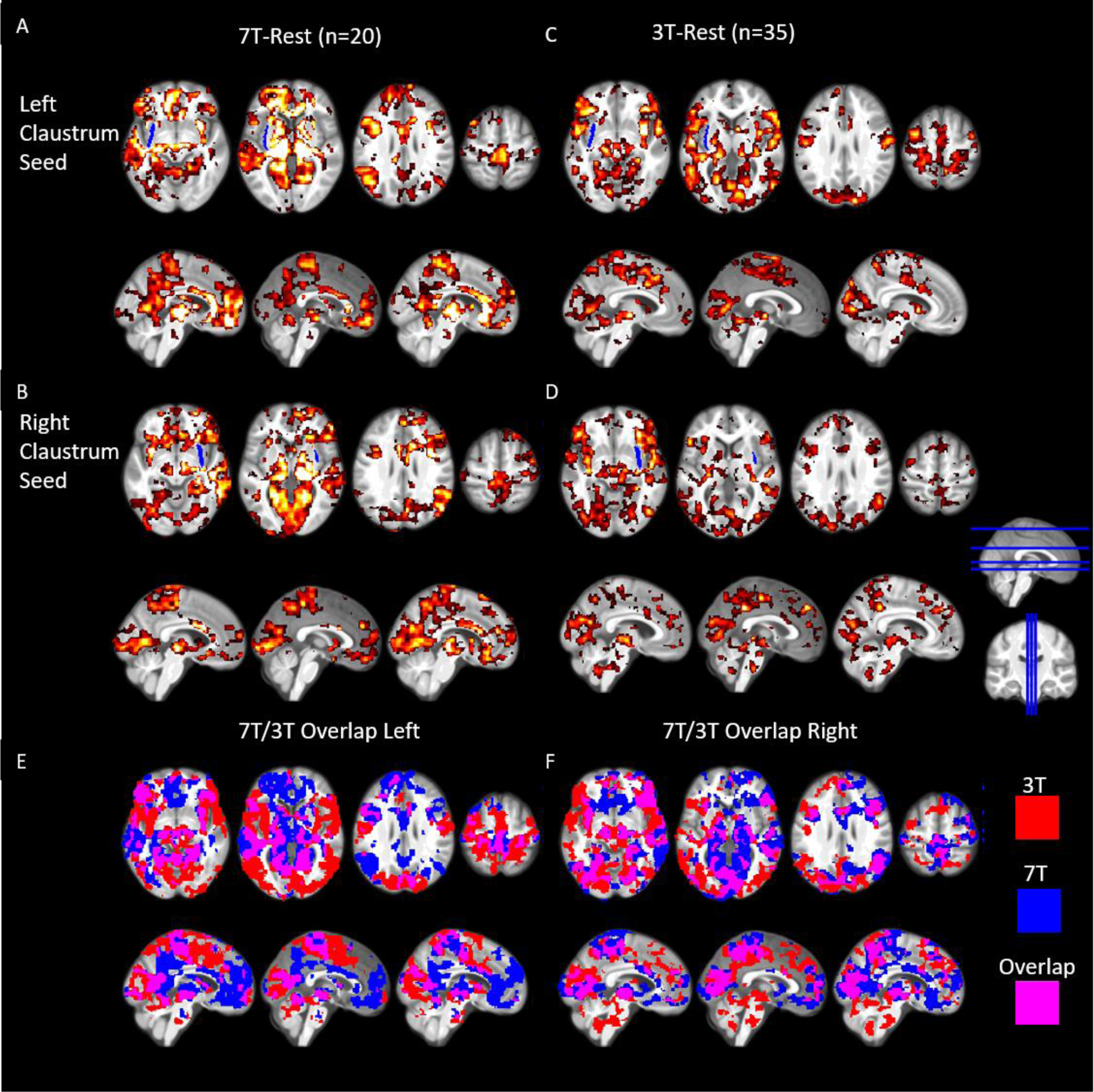
Resting state connectivity of the left and right claustrum at 7T and 3T showing widespread connectivity to cortical and subcortical regions. A) RSFC of the left claustrum in *7T-Rest.* B) RSFC of the right claustrum in *7T-Rest.* C) RSFC of the left claustrum in *31-Rest.* D) RSFC of the right claustrum in *3T-Data.* E) and F) show the overlap of these thresholded RSFC maps. Data were voxelwisethresholded at p<0.001 followed by FWE cluster correction. Blue ROI in A-D represents an average claustrum dilated by 1.5 mm (for visualization only). Cl=claustrum.

We found a more extensive pattern of claustrum functional connectivity in *3T-Rest,* however, the regions displaying FC with claustrum were largely similar to *7T-Rest.* In *3T-Rest* we identified connectivity with: the thalamus, mainly the pulvinar; nucleus accumbens; visual cortex; both anterior and posterior cingulate cortex; PFC, precuneus; angular gyrus; sensorimotor cortex; parahippocampal gyrus; temporal gyri; and IPS (Fig 2C and D and Table S3 and S4). The similar pattern of claustrum functional connectivity across *7T-Rest* and *3T-Rest,* and the bilateral nature of claustrum functional connectivity (i.e. left claustrum functionally connects with right) suggest our confound-corrected claustrum timeseries is not artifactual.

### Extrinsic and Default Mode Network overlap with claustrum functional connectivity

Given the high degree of claustrum functional connectivity with regions involved in cognitive control (e.g. ACC) and prior literature showing claustrum involvement in top-down cognitive processing (White et al., 2018), we sought to quantify the overlap between resting-state claustrum connectivity, the cognitive conflict task positive network (i.e. extrinsic mode network; EMN) (Hugdahl., 2015) and the task negative network (i.e. default mode network; DMN) (Raichle et al., 2001) evoked by the multi-source interference task (MSIT; Bush et al., 2003). MSIT features a difficult condition where there is conflict between the position and identity of a number that must be selected and an easy condition, where there is no conflict. The neural activation associated with these conditions are contrasted to a motor and visual control condition. Examining the difficult-block greater than easy-block contrast identified the standard EMN, including fronto-parietal network (FPN), dorsal-attention network (DAN), and dorsal ACC (Fox et al., 2005; see Fig 3A and Table S5). The easy-block greater than difficult-block contrast identified the DMN. We found left and right claustrum functional connectivity overlapped with more than half of the regions in both the extrinsic mode and default mode networks (Fig 3C). Additionally, every major region with significant task activation was also functionally connected to the claustrum (i.e. at least partially overlapped).

**Fig 3.**
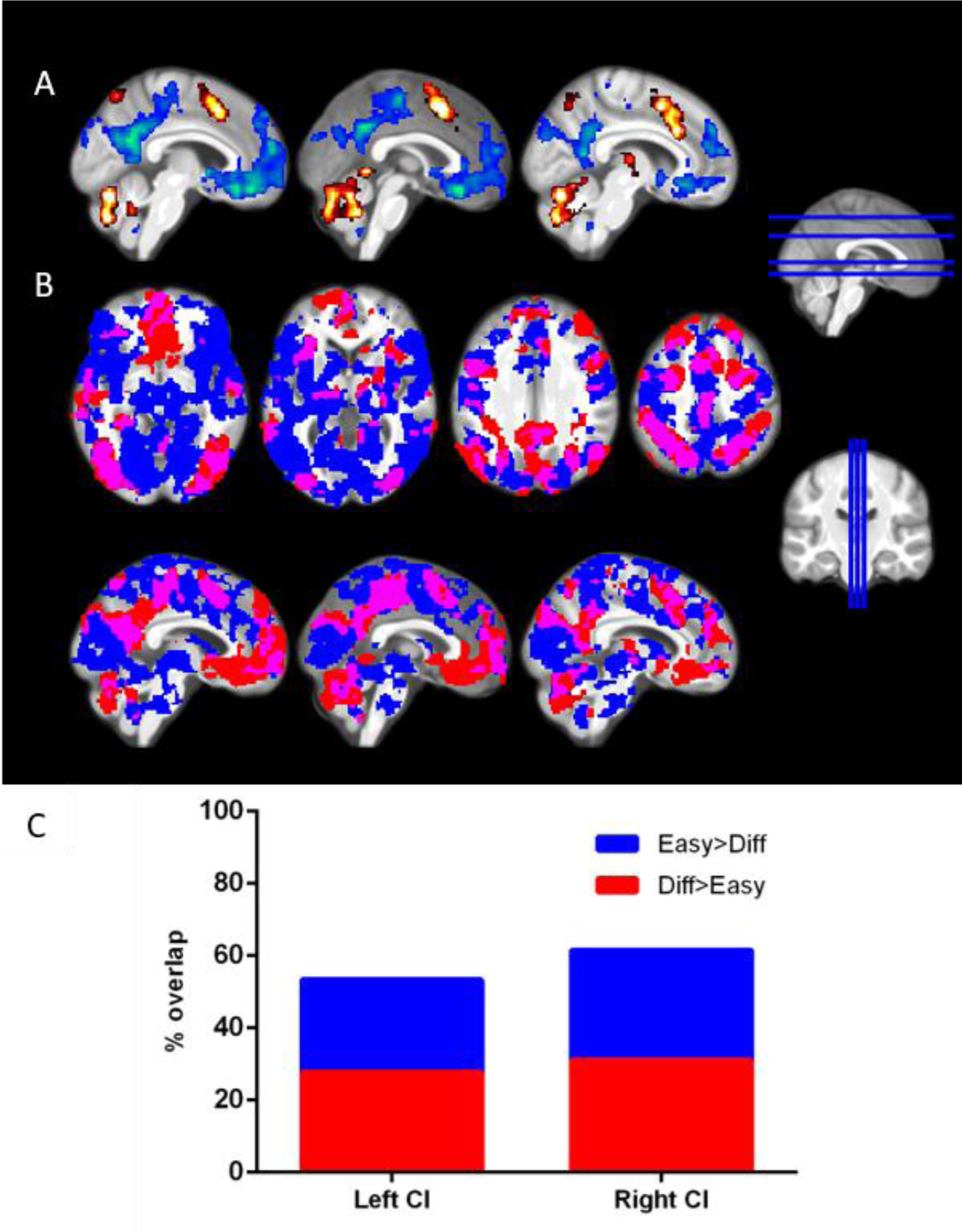
Claustrum functional connectivity substantially overlaps with DMN and EMN. A) Contrasting the difficult block and the easy block of MSIT reveals DMN (easy > difficult, shown in blue) and EMN (difficult > easy, shown in red). B) Areas functionally connected with L/R claustrum in 3T-Rest (blue) and the DMN and EMN shown above (red) show a high degree of overlap (purple) C) Quantification of DMN/EMN voxels from panel A and the percent that also have FC with claustrum. Claustrum FC maps are the same as shown in Figure 2B. Task contrasts maps used a p< 0.001 threshold with FWE cluster correction. Easy=easy condition of MSIT, Diff=Difficult condition of MSIT, Cl=claustrum, DMN=default mode network, EMN=extrinsic mode network.

### MSIT performance

Consistent with past literature, we found a main effect of reaction time (RT; Fu6,56.38=374.2, p<0.05), with RT for tapping<easy<difficult trials within the MSIT, and all trial types differed from one another in terms of RT (Fig 4A).

**Fig 4.**
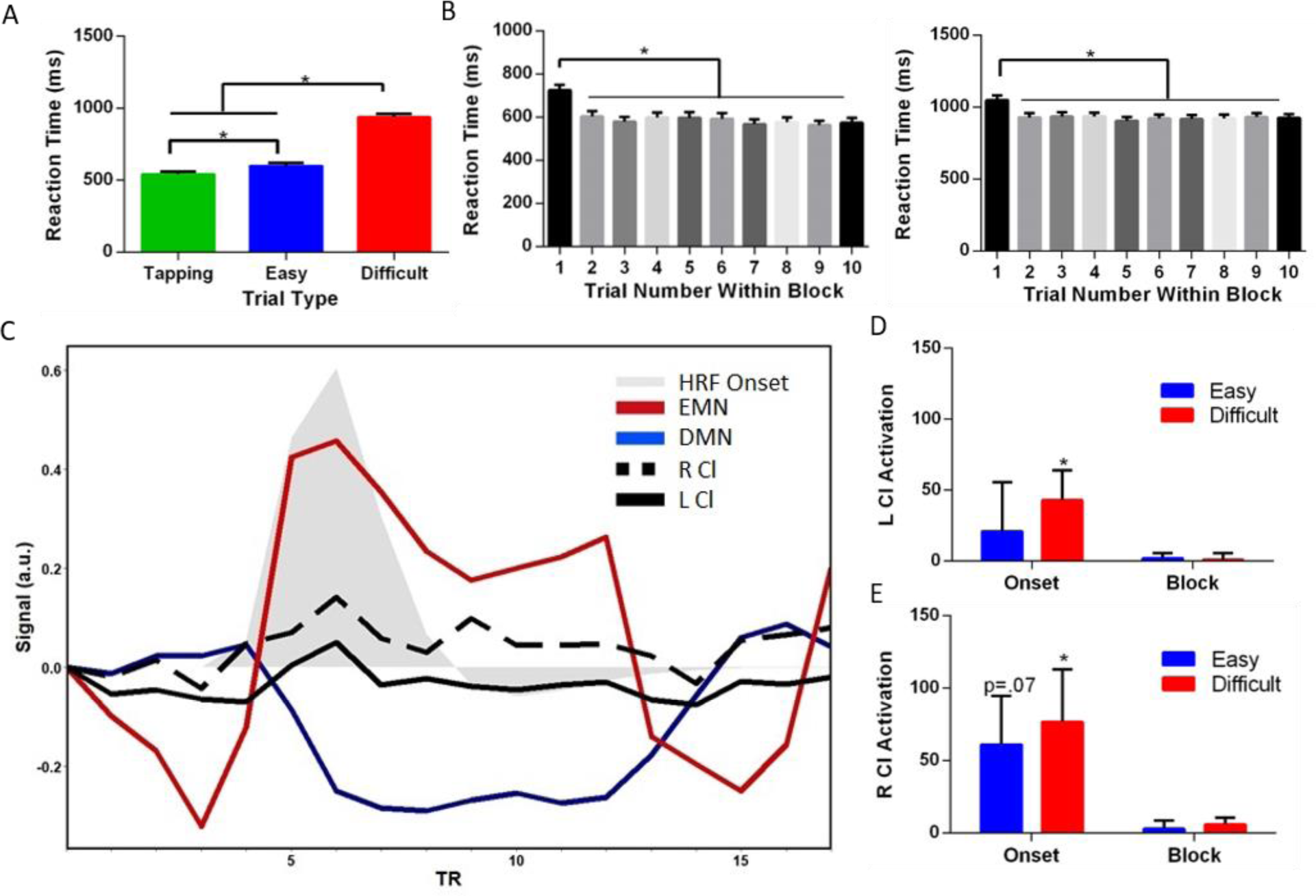
Claustrum activates when cognitive load peaks. A) Reaction time performance in MSIT varies based on the condition, with the difficult blocks having the slowest reaction times. B) The first trial of the easy and difficult conditions is behaviorally unique from the other trials in the block. As a result, we modelled the onset separately from the remainder of the block for difficult and easy conditions in an adaptive model to capture individual variability. C) Time course of claustrum shows a response during the difficult task, but exclusively during the onset of that task. The contrast map localizing DMN and the contrast map localizing EMN (Fig 3A) were used as regions of interest. We extracted the signal from these areas along with the left and right claustrum averaged over every difficult condition for all subjects. The task starts at the third TR and ends at the 11^th^. TRs 1:2 and 12:15 show the control condition. We under laid a canonical hemodynamic response function (HRF) convolved with an onset of duration 3.2 second (two trials in duration). D) Left and E) right claustrum activates at the onset – but not block – of the difficult MSIT condition. Error bars show standard error of the mean. * represents a p value of <0.05. HRF=Hemodynamic Response Function, Cl=claustrum, DMN=default mode network, EMN=extrinsic mode network.

When analyzing RT at the group level, we observed that RT was significantly higher for the first trial of easy and difficult blocks than the remaining 9 trials, a finding observed in many cognitive tasks (Blais et a., 2014) (Fig 4B) (Difficult: F6.27,200.7=44.17, p<0.05; Easy: F4.33,138.5=25.43, p<0.05). These data suggest that after the first trial participants transition into a stable level of performance and therefore the beginning or onset of difficult and easy conditions should be modelled separately from the block (creating 5 conditions to be modelled: tapping, easy onset, easy block, difficult onset, difficult block). While at the group level a transition from high initial RT for the first trial to a near-mean RT by the second trial occurred, we observed variability within subjects and found that subjects sometimes took longer to transition into this stable level of performance where RT was close to the block mean. That is, for a given block of stimuli some subjects had clearly slower RTs for both the first and second/third trials relative to the RT mean. In order to capture this inter-and intra-individual variation as well as possible, we allowed the duration of the easy onset and difficult onset to vary based on RT of the subject. For start time of onset events, we used the beginning of the task block. For duration, we created an adaptive program that defined the duration of the easy onset and difficult onset events as either the first trial RT, the second trial RT (+ trial one duration of 1.6 s), or the third trial RT (+ trial one and two duration of 3.2 s). The choice of which trial RT to use as duration (one, two, or three) was based on the z scores for each of these trials using the mean calculated for each subject of the same task type (difficult or easy) for the fourth to tenth trials over all runs. Specifically, trial one RT was the onset duration if trial two had a z-score ≤ to 0.5; trail two was used if trail one RT had a z score > 0.5 and if trial three had a z score ≤ to 0.5; trial three was used if these prior stipulations were not met. Hence, the onset event duration was adapted based on individual level behavioral data, which ensured that for each subject we captured the period of greatest cognitive conflict, reflective of the switch to a new task. The block of the task type was modelled as the remainder of the block not included in onset. We used this hybrid event-block design for all task analysis of MSIT and found activations and deactivations for the onset and block for both easy and difficult conditions across the brain (Table S5, S6, S7, and S8).

### Claustrum responds to the switch to task onset

The onset of tasks involved switching from performing a tapping visuomotor control task to performing either a no-cognitive conflict easy task or a cognitive conflict difficult task. When analyzing claustrum activation independent of surrounding insula and putamen using this RT adaptive model, we found that claustrum significantly responded to difficult onset (Left: t_32_=2.1, p<0.05; Right: t_32_=2.2, p<0.05), though the same effect was not found for easy onset (Left: t_32_=0.6, p>.2; Right: t_32_=1.8, p=0.07, Fig 4D-E). However, we found no evidence that claustrum responded to the difficult block nor to the easy block and found no evidence that claustrum responded to difficult onset more than to easy onset (all p>0.2). This strongly suggests that claustrum is involved in the switch to active cognitive control, rather than maintaining performance during high cognitive demand.

## Discussion

In this study, we provide a novel approach we term Small Region Confound Correction to detect the activity and functional connectivity of the human claustrum. In doing so, we find that the claustrum is strongly functionally connected to cingulate and prefrontal cortices at rest and that there is considerable overlap between claustru|m connectivity maps and cognitive task-related networks. Supporting a role of the claustrum in cognition, we show that claustrum is activated at the onset of a demanding cognitive task, which is also associated with the onset of EMN engagement.

We isolated a distinct claustrum signal by regressing putamen and insula signals from the claustrum. This is a conservative approach, assuming that any linear relationship between claustrum and insula/putamen is a result of a partial volume effect, defined as signal from outside structures being erroneously incorporated into an ROI by virtue of the ROI only occupying a portion of a measured volume (Dukart and Bertolino, 2014; Du et al., 2014). Thus, while our approach mitigates the partial volume effect, a degree of true claustrum signal may be dampened. However, our resting state analyses showing similar RSFC across left/right claustrum, similar RSFC over two datasets, bilateral claustrum RSFC, and task response of the claustrum, argue that the signal that we do derive is sufficiently robust. This approach offers a generalizable method to assay the function of the claustrum, or other small or oddly shaped neural structures, independent of partial volume effects from surrounding structures. SRCC serves as a platform for studying a host of regions across the brain, such as the habenula (Shelton et al., 2012; Hetu et al., 2016) and other thalamic association nuclei.

Our claustrum functional connectivity data reveals co-activation of claustrum with executive cortical regions including ACC and medial PFC. This is in line with decades of neuronal tract tracing studies from rodent to monkeys (for review see Mathur, 2014), and particularly in line with reports in rat indicating dense claustrum connections with ACC and prelimbic PFC (Smith and Alloway, 2010; White et al., 2017) and in with area 24 in the common marmoset (Reser et al., 2017). The present work also indicates that claustrum is functionally connected to posterior cingulate cortex, precuneus, angular gyrus, cuneus, visual cortex, and sensorimotor cortex, which is in line with connections from claustrum to parietal association cortex and sensorimotor cortices in rats (White et al., 2017), connections from claustrum to visual cortex in cats (LeVay and Sherk, 1981) and projections from claustrum to parietal cortex in monkeys (Gamberini et al., 2017). The more extensive pattern of claustrum functional connectivity observed in *3T-Rest* may be a result of the larger sample size compared to *7T-Rest* (35 for 3T vs. 20 for 7T). The additional functional connectivity seen in *3T-Rest* with midbrain and nucleus accumbens in particular, is likely to reflect indirect connectivity as direct connections between these regions are not currently strongly supported in the tract tracing literature. Alternatively, functional connectivity of claustrum with these structures may reflect connections that are unique to the human brain.

Our data also suggest involvement of the claustrum at the start of a difficult condition when new rules come into play that require a change of cognitive strategy and task set instantiation (Dosenbach et al., 2006). Significant claustrum activation was only observed at the switch from the stimulus-response-based tapping task to the difficult condition of MSIT. As subjects transitioned to this new rule set, average reaction time decreased significantly compared with later trials in the task block. This transition also was met with an emergence of the task positive/extrinsic mode network. A possible interpretation of these data could be that the claustrum is involved in action inhibition as cognitive demand soars. However, neither optogenetic inhibition nor activation of axon terminals of a major excitatory input source to the claustrum, the ACC, affects motor activity in mice (White et al., 2018). Our results cannot be explained through alterations in sensory binding or motor processes, which are both proposed roles for the claustrum, as we observed claustrum activation when controlling for sensory input and motor responses. The data also do not suggest a role for claustrum in resolving cognitive conflict, as claustrum did not show sustained activation during the block of the difficult condition, whereas the EMN did (Fig 3A).

In summary, we showed that even at 1.5mm spatial resolution attained with 7T fMRI, the claustrum BOLD signal bears unsettling similarity with the insula and putamen, and an additional processing step of SRCC allowed us to isolate a claustrum signal independent of the surrounding regions. Using this method we find that in in the human – like in other species – the claustrum has wide-ranging cortical connectivity, including default and extrinsic mode networks, to sensory regions. Additionally, we show that claustrum activity peaks when switching to a cognitive conflict task and that this activity could not be explained by changes in consciousness or sensorimotor processing. These data broadly support a role of the claustrum in cognitive control and are consistent with recent studies in mice (White et al., 2017).

## Acknowledgments

This work was supported by National Institute on Alcohol Abuse and Alcoholism grants K22AA021414, R01AA024845 (B.N.M.) and F31AA024683 (M.H.P.), Whitehall Foundation grant 2014-12-68 (B.N.M.), National Institute of General Medical Sciences grant T32008181 (M.G.W.), National Institute of Neurological Disorders and Stroke grant T32NS063391 (M.G.W.), National Institute on Drug Abuse grant R03DA042336 (F.S.B.), and National Center for Complementary and Integrative Health grant R01AT007176 (D.A.S.); The authors also are grateful for the assistance of Dr. Rao Gullapalli and the Core for Translational Research in Imaging @ Maryland (C-TRIM) and the Center for Metabolic Imaging and Therapeutics (CMIT).

## Author contributions

DAS and BNM conceived of the research. DAS, SK, MGW, and BNM designed research. MP generated claustrum masks. DAS, SK, FSB, MGW and BNM analyzed data. DAS, SK, and BNM wrote the manuscript.

## References

1. Behzadi, Y., Restom, K., Liau, J., & Liu, T. T. (2007). A component based noise correction method (CompCor) for BOLD and perfusion based fMRI. Neuroimage, 37(1), 90–101.

2. Blais, C., Stefanidi, A., & Brewer, G. A. (2014). The gratton effect remains after controlling for contingencies and stimulus repetitions. Frontiers in Psychology, 5, 1207.

3. Bush, G., Shin, L., Holmes, J., Rosen, B., & Vogt, B. (2003). The multi-source interference task: Validation study with fMRI in individual subjects. Molecular Psychiatry, 8(1), 60.

4. Crick, F. C., & Koch, C. (2005). What is the function of the claustrum? Philosophical Transactions of the Royal Society of London. Series B, Biological Sciences, 360(1458), 1271–1279.

5. Dosenbach, N. U., Visscher, K. M., Palmer, E. D., Miezin, F. M., Wenger, K. K., Kang, H. C., Burgund, D., Grimes, A. L., Schlaggar, B. L., & Petersen, S. E. (2006). A core system for the implementation of task sets. Neuron, 50(5), 799–812.

6. Du, Y. P., Chu, R., & Tregellas, J. R. (2014). Enhancing the detection of BOLD signal in fMRI by reducing the partial volume effect. Computational and Mathematical Methods in Medicine, 2014, 973–972.

7. Dukart, J., & Bertolino, A. (2014). When structure affects function-the need for partial volume effect correction in functional and resting state magnetic resonance imaging studies. PloS One, 9(12), e114227.

8. Edelstein, L., & Denaro, F. (2004). The claustrum: A historical review of its anatomy, physiology, cytochemistry and functional significance. Cellular and molecular biology, 50: 675–702.

9. Fox, M. D., Snyder, A. Z., Vincent, J. L., Corbetta, M., Van Essen, D. C., & Raichle, M. E. (2005). The human brain is intrinsically organized into dynamic, anticorrelated functional networks. Proceedings of the National Academy of Sciences of the United States of America, 102(27), 9673–9678.

10. Gamberini, M., Passarelli, L., Bakola, S., Impieri, D., Fattori, P., Rosa, M. G., & Galletti, C. (2017). Claustral afferents of superior parietal areas PEc and PE in the macaque. Journal of Comparative Neurology, 525(6), 1475–1488.

11. Gorgolewski, K. J., Mendes, N., Wilfling, D., Wladimirow, E., Gauthier, C. J., Bonnen, T., Cozatl, R. (2015). A high resolution 7-tesla resting-state fMRI test-retest dataset with cognitive and physiological measures. Scientific Data, 2, 140054.

12. Hetu, S., Luo, Y., Saez, I., D'ardenne, K., Lohrenz, T., & Montague, P. R. (2016). Asymmetry in functional connectivity of the human habenula revealed by high-resolution cardiac-gated resting state imaging. Human Brain Mapping, 37 (7), 2602–2615.

13. Hugdahl, K., Raichle, M. E., Mitra, A., & Specht, K. (2015). On the existence of a generalized non-specific task-dependent network. Frontiers in Human Neuroscience, 9, 430.

14. LeVay, S., & Sherk, H. (1981). The visual claustrum of the cat. I. structure and connections. The Journal of Neuroscience, 1 (9), 956–980.

15. Mathur, B. N. (2014). The claustrum in review. Frontiers in Systems Neuroscience, 8, 48.

16. Mathur, B. N., Caprioli, R. M., & Deutch, A. Y. (2009). Proteomic analysis illuminates a 495 novel structural definition of the claustrum and insula. Cerebral Cortex, 19 (10), 2372–2379.

17. Murphy, K., & Fox, M. D. (2017). Towards a consensus regarding global signal regression for resting state functional connectivity MRI. Neuroimage, 154, 169–173.

18. Muschelli, J., Nebel, M. B., Caffo, B. S., Barber, A. D., Pekar, J. J., & Mostofsky, S. H. (2014). Reduction of motion-related artifacts in resting state fMRI using aCompCor. Neuroimage, 96, 22–35.

19. Nakashima, M., Uemura, M., Yasui, K., Ozaki, H. S., Tabata, S., & Taen, A. (2000). An anterograde and retrograde tract-tracing study on the projections from the thalamic gustatory area in the rat: Distribution of neurons projecting to the insular cortex and amygdaloid complex. Neuroscience Research, 36 (4), 297–309.

20. Pan, W. X., Mao, T., & Dudman, J. T. (2010). Inputs to the dorsal striatum of the mouse reflect the parallel circuit architecture of the forebrain. Frontiers in Neuroanatomy, 4, 147.

21. Patel, A. X., Kundu, P., Rubinov, M., Jones, P. S., Vértes, P. E., Ersche, K. D., Suckling, J., Bullmore, E. T. (2014). A wavelet method for modeling and despiking motion artifacts from resting-state fMRI time series. Neuroimage, 95, 287–304.

22. Power, J. D., Plitt, M., Laumann, T. O., & Martin, A. (2017). Sources and implications of whole-brain fMRI signals in humans. Neuroimage, 146, 609–625.

23. Raichle, M. E., MacLeod, A. M., Snyder, A. Z., Powers, W. J., Gusnard, D. A., & Shulman, G. L. (2001). A default mode of brain function. Proceedings of the National Academy of Sciences of the United States of America, 98 (2), 676–682.

24. Reser, D. H., Majka, P., Snell, S., Chan, J. M., Watkins, K., Worthy, K., Quiroga, M. D., & Rosa, M. G. (2017). Topography of claustrum and insula projections to medial prefrontal and anterior cingulate cortices of the common marmoset (Callithrix jacchus). Journal of Comparative Neurology, 525 (6), 1421–1441.

25. Sato, F., Akhter, F., Haque, T., Kato, T., Takeda, R., Nagase, Y., Yoshida, A. (2013). Projections from the insular cortex to pain-receptive trigeminal caudal subnucleus (medullary dorsal horn) and other lower brainstem areas in rats. Neuroscience, 233, 9–27.

26. Seminowicz, D. A., & Davis, K. D. (2007). Pain enhances functional connectivity of a brain network evoked by performance of a cognitive task. Journal of Neurophysiology, 97 (5), 3651–3659.

27. Seminowicz, D. A., Wideman, T. H., Naso, L., Hatami-Khoroushahi, Z., Fallatah, S., Ware, M. A., Stone, L. S. (2011). Effective treatment of chronic low back pain in humans reverses abnormal brain anatomy and function. The Journal of Neuroscience, 31 (20), 7540–7550. doi:10.1523/JNEUROSCI.5280–10.2011 [doi]

28. Shelton, L., Pendse, G., Maleki, N., Moulton, E. A., Lebel, A., Becerra, L., & Borsook, D. (2012). Mapping pain activation and connectivity of the human habenula. Journal of Neurophysiology, 107(10), 2633–2648.

29. Smith, J. B., & Alloway, K. D. (2010). Functional specificity of claustrum connections in the rat: Interhemispheric communication between specific parts of motor cortex. The Journal of Neuroscience, 30(50), 16832–16844.

30. Smith, J. B., Radhakrishnan, H., & Alloway, K. D. (2012). Rat claustrum coordinates but does not integrate somatosensory and motor cortical information. Journal of Neuroscience, 32(25), 8583–8588.

31. Smith, S. M., Beckmann, C. F., Andersson, J., Auerbach, E. J., Bijsterbosch, J., Douaud, G., Harms, M. P. (2013). Resting-state fMRI in the human connectome project. Neuroimage, 80, 144–168.

32. Torgerson, C. M., Irimia, A., Goh, S., & Van Horn, J. D. (2015). The DTI connectivity of the human claustrum. Human Brain Mapping, 36(3), 827–838.

33. Wang, Q., Ng, L., Harris, J. A., Feng, D., Li, Y., Royall, J. J., Koch, C. (2017). Organization of the connections between claustrum and cortex in the mouse. Journal of Comparative Neurology, 525(6), 1317–1346.

34. White, M. G., & Mathur, B. N. (2018). Frontal cortical control of posterior sensory and association cortices through the claustrum. Brain Structure and Function, 1–8.

35. White, M. G., Cody, P. A., Bubser, M., Wang, H., Deutch, A. Y., & Mathur, B. N. (2017). Cortical hierarchy governs rat claustrocortical circuit organization. Journal of Comparative Neurology, 525(6), 1347–1362.

36. White, M. G., Panicker, M., Mu, C., Carter, A. M., Roberts, B. M., Dharmasri, P. A., & Mathur, B. N. (2018). Anterior cingulate cortex input to the claustrum is required for top-down action control. Cell Reports, 22(1), 84–95.

37. Woo, C., Krishnan, A., & Wager, T. D. (2014). Cluster-extent based thresholding in fMRI analyses: Pitfalls and recommendations. Neuroimage, 91, 412–419.

